# Measures of states of consciousness during attentional and cognitive load

**DOI:** 10.1101/586149

**Authors:** André S. Nilsen, Bjørn E. Juel, Johan F. Storm

## Abstract

**Background:** Developing and testing methods for reliably assessing states of consciousness in humans is important for both basic research and clinical purposes. Several potential measures, partly grounded in theoretical developments, have been proposed, and some of them seem to reliably distinguish between conscious and unconscious brain states. However, the degrees to which these measures may also be affected by changes in brain activity or conditions that can occur within conscious brain states have rarely been tested. In this study we test whether several of these measures are modulated by attentional load and related use of cognitive resources.

**Methods:** We recorded EEG from 12 participants while they passively received three types of stimuli: (1) transcranial magnetic stimulation (TMS) pulses (for measuring perturbational complexity), (2) auditory stimuli (for detection of auditory pattern deviants), or (3) audible clicks from a clock (spontaneous EEG, for measures of signal diversity and functional connectivity). We investigated whether the measures significantly differed between the passive condition and a attentional and cognitively demanding working memory task.

**Results:** Our results showed that in the attention-based auditory P3b ERP measure (global auditory pattern deviant) was significantly affected by increased attentional and cognitive load, while the various measures based on spontaneous and perturbed EEG were not affected.

**Conclusion:** Measures of conscious state based on complexity, diversity, and effective connectivity, are not affected by attentional and cognitive load, suggesting that these measures can be used to test both for the presence and absence of consciousness.

## Introduction

The ability to objectively detect the presence, absence, level, or degree of consciousness in individuals is of both clinical and theoretical relevance. Here, we focus on the most basic concept of consciousness, defined as phenomenological experience, or what it feels like for a person (or in general: a system) to be in a state, for example for a human to be happy. Thus, if an individual or a system has any phenomenological experience at all, it is considered to be conscious. It is generally assumed that humans are conscious when they report being conscious, and they are considered to be largely unconscious during dreamless states of sleep, general anesthesia, or during coma. Several methods aiming at objectively assessing or quantifying the presence or absence of conscious or unconscious states, or the level or degree of consciousness, have been developed, based on various metrics such as behavior (i.e. the Glasgow Coma Scale; (Teasdale & Jennett, 1974), brain metabolism (i.e. brain glucose consumption; (Shulman, Hyder, & Rothman, 2009), and electrophysiological activity (i.e. the electroencephalogram (EEG)-based BIS index; (Myles, Leslie, McNeil, Forbes, & Chan, 2004). While several measures have shown promising results in being able to reliably distinguish awake states from states of general anesthesia or sleep, it’s important to investigate whether and how they may change within each of these states. For example, it’s important that a clinical marker is not strongly affected by attention and other cognitive functions, which both can vary during normal wakefulness, and can be impaired in some groups of patients.

Two of the EEG-based measures inspired by theories of consciousness are the perturbational complexity index (PCI; (Casali et al., 2013) and the P3b component of the sensory event related potentials (ERP) elicited by the so-called “global-local” oddball paradigm of sensory stimulation (Bekinschtein et al., 2009; King et al., 2013; Sergent, Baillet, & Dehaene, 2005). The PCI is a measure based on quantifying the spatiotemporal complexity of the global EEG responses to local transcranial magnetic stimulation (TMS), and is inspired by the integrated information theory of consciousness (Oizumi, Albantakis, & Tononi, 2014). The P3b ERP is a late EEG response reflecting detection of temporal pattern irregularities, usually elicited by the “auditory global-local” paradigm, and is inspired by the global neuronal workspace (GNW) theory of consciousness (Baars, 1997; Dehaene, Kerszberg, & Changeux, 1998; Dehaene & Naccache, 2001). However, in the original studies employing the auditory global local paradigm to elicit the P3b response, it was observed that participants doing a distracting task (or who were just allowed to mind-wander) during the paradigm, did not show the same response as those that specifically paid attention to the auditory stimuli (Bekinschtein et al., 2009; King et al., 2013). The authors argued that the P3b can be more correctly described as a marker of being conscious of specific content rather than being in a generally conscious state. Thus, the presence of the P3b wave may be a strong indicator of conscious access or attention to a stimulus; but a lack of the P3b wave might only indicate faulty sensory processing, failure to follow commands, or internally generated conscious states like dreaming, and not indicate overall state unconsciousness per se. In contrast, according to several studies, the PCI is apparently not dependent on or noticeably influenced by attending to any specific stimulus (Casali et al., 2013; Casarotto et al., 2016). However, the TMS method gives not only purely magnetic stimulations of the cortex, but is normally also accompanied by a high-frequency audible “click” as well as a noticeable somatosensory sensation caused by stimulation of nerves in the scalp. While both sensations can be reduced by using noise masking and foam padding, respectively, they may still be perceived and are time-locked to the magnetic stimulation. Therefore, it is conceivable that the EEG-responses to TMS may be contaminated by sensory responses to these TMS-related auditory and somatosensory stimuli. There is currently disagreement to what extent such sensory contamination may occur (Biabani, Fornito, Mutanen, Morrow, & Rogasch, 2019; Conde et al., 2019), but if it’s significant, it’s possible that sufficient attentional distractions (such as concurrent task performance) might have similar effects on the TMS-evoked EEG responses as have been seen for the P3b. However, the PCI measure has shown similar values in the dreaming state, when one is presumably unconscious of the TMS stimulations, as in the awake state (Casarotto et al., 2016). Therefore, it seems improbable that attention towards sensory aspects of TMS stimulation affects PCI, when the TMS-EEG procedure is properly executed (Belardinelli et al., 2019). However, there is an alternative hypothesis: the observed difference in response to auditory pattern irregularities (P3b) might not only be modulated by attentional focus but also by the cognitive load and the use of attentional resources demanded by the task itself.

Both attentional and cognitive load can alter the underlying neural activity measured by EEG. Focusing on different sensory modalities are linked to increased activations in the related primary and secondary sensory areas (Pessoa, Kastner, & Ungerleider, 2003), as well as different sensory related functional networks (Smith et al., 2009), while task engagement is linked to variations in power spectral density distributions in EEG (Hsieh, Ekstrom, & Ranganath, 2011), general task related networks (Fox et al., 2005), as well as task specific modulation of activity in specific brain regions. Such changes may contaminate measures of conscious and non-conscious states if they are sensitive to properties not specifically relevant for consciousness. Therefore, it’s important to verify that any measure of consciousness is at least independent of attentional focus and cognitive load.

This study aimed to investigate the effect of attentional and cognitive load modulation on PCI, P3b, and some other recently developed EEG-based measures that have shown promise for classification of states of consciousness; Lempel Ziv complexity (LZc; Schartner, Pigorini, et al., 2017), synchrony coalition entropy (SCE; Schartner et al., 2015) and amplitude coalition entropy (ACE; Schartner et al., 2015). In addition, we implemented a novel measure, based on directed functional connectivity estimated using the Directed Transfer Function (DTF; Kamiński & Blinowska, 1991), that has recently shown promise in separating the conscious from the non-conscious state during anesthesia, with high time resolution (Juel, Bremnes, et al., 2018; Juel, Romundstad, Kolstad, Storm, & Larsson, 2018). To test whether attentional and cognitive load modulation affect these six measures, we implemented a within subject design with a low-demand cognitive task related to the methodology behind the measures, and an high-demand adaptive working memory task aimed at constantly loading participants’ attentional and cognitive resources. Based on the difference in nature of the different measures and prior findings, we hypothesized that none of the measures should be detrimentally affected by attentional and cognitive modulation.

## Methods

### Participants

12 healthy participants were recruited through written adverts and posters at the University of Oslo and Oslo University Hospital (*n*_*female*_=7, age range 21-55 yrs, median 27.5 yrs). All participants received oral and written information and signed informed consent prior to the study. All participants could withdraw their consent at any point during the study, and received monetary compensation regardless of whether they completed the study or not. Inclusion criteria were age between 18 and 60 years, no magnetic resonance imaging (MRI) contraindications, no TMS-EEG risk factors (Rossi, Hallett, Rossini, Pascual-Leone, & Safety of TMS Consensus Group, 2009), easily observable motor evoked potentials, and low level of muscular activations in response to TMS stimulation around target areas (BA6, BA7). The study was approved by the Regional Committees for Medical and Health Research Ethics (REC South-East Norway: 2014/1952-1).

### Measures & Stimuli

#### Spontaneous measures

For measures based on spontaneous EEG we aimed to calculate signal complexity (LZc) and entropy (ACE, SCE), as well as directed functional connectivity (DTF). To measure signal complexity we measured the discretized EEG signal’s compressibility through Lempel Ziv compression using the 1976 variant (Lempel & Ziv, 1976) with algorithmic implementation by Kaspar and Schuster (1987).

To measure entropy we used ACE and SCE (Schartner, Pigorini, et al., 2017; Schartner et al., 2015). ACE is a variant of the measure implemented by Shanahan (2010), and is measured on discrete spontaneous EEG data (like LZc) by calculating the entropy of the distribution of channel coalitions over time, and normalized with respect to the entropy of a shuffled version of the EEG data. SCE is similar to the ACE, except that coalitions over channels are determined by whether pairs of channels are phase synchronized or not.

LZc, ACE, and SCE, have previously been used to separate conscious and nonconscious states, as well as distinguishing between the normal and psychedelic conscious state (Fan, Yeh, Chen, Shieh, & Others, 2011; Ferenets et al., 2006; Hudetz, Liu, Pillay, Boly, & Tononi, 2016; Schartner, Carhart-Harris, Barrett, Seth, & Muthukumaraswamy, 2017; Schartner, Pigorini, et al., 2017; Schartner et al., 2015).

To measure directed functional connectivity (i.e. effective connectivity; (Sporns, 2007), we calculated DTF applied directly to spontaneous EEG data (Kamiński & Blinowska, 1991). DTF quantifies the information carried by any one time-series about any other (including itself) given a multivariate autoregressive (MVAR) model (Greenblatt, Pflieger, & Ossadtchi, 2012), and has shown promise in distinguishing between states of consciousness (De Gennaro et al., 2005; Höller et al., 2014). Recently, we used the DTF to successfully distinguish between states of wakefulness and several different types of general anesthesia using only raw EEG (Juel, Bremnes, et al., 2018; Juel, Romundstad, et al., 2018), providing further evidence that the DTF is sensitive to changes in the state of consciousness.

#### Evoked measures

For measures based on ERP and TMS-evoked potential (TEP) data, we focused on P3b and PCI, respectively. PCI is based on methods inspired by the integrated information theory of consciousness (Casali et al., 2013; Ferrarelli et al., 2010; Marcello Massimini et al., 2005; M. Massimini, Tononi, & Huber, 2009), and requires ∼100-300 single pulses of TMS repeated at a low rate (<1 Hz) to locally perturb the cortex, combined with high density EEG to measure the response (TEP). The TEPs (averaged across trials) are then binarized at the source level and the overall complexity of the set of responses is calculated using LZc. By assessing the the overall complexity of the responses, PCI depends both on integration and differentiation within the cortex, and has so far shown near 100% accuracy in distinguishing between conscious states vs. non-conscious states, as compared with subjective reports and/or behavioral markers, under a variety of conditions (Casarotto et al., 2016).

The P3b marker elicited by the global local paradigm measures the ERP of the late (>300 ms) EEG response to a break in global auditory patterns relative to habituated patterns (see (Bekinschtein et al., 2009)). The auditory paradigm employed to elicit the P3b component of the ERP was similar to that of King et al (2013). Specifically, we presented 8 blocks of 105-135 trials. One trial consisted of five 50ms tones with 150ms inter-tone interval, with a 1150ms silent inter trial interval. The first four tones were identical with the fifth deviating or not, depending on trial condition. A trial could be local deviant which was defined as the fifth tone being higher or lower in frequency than the first four. The first 15 trials within a block were identical, and either local deviant or not. This habituated a global pattern of local deviance. Global deviance was defined as a change in frequency of the fifth tone compared to the fifth tone in the majority (80%) of trials within a block (also the habitutuated trials). This resulted in four trial conditions; 1) local and global non-deviant, 2) local non-deviant and global deviant, 3) local deviant and global non-deviant, or 4) local and global deviant. In total, we presented 8 blocks (2 of each condition) with 20-30 global deviants (90-120 trials, total 30-40 minutes). The trials were semi-randomized so that no two trials with 20% probability appeared after each other, and with a minimum of two 80% probability trials in between two 20% probability trials. See Figure 1.

**Figure 1.**
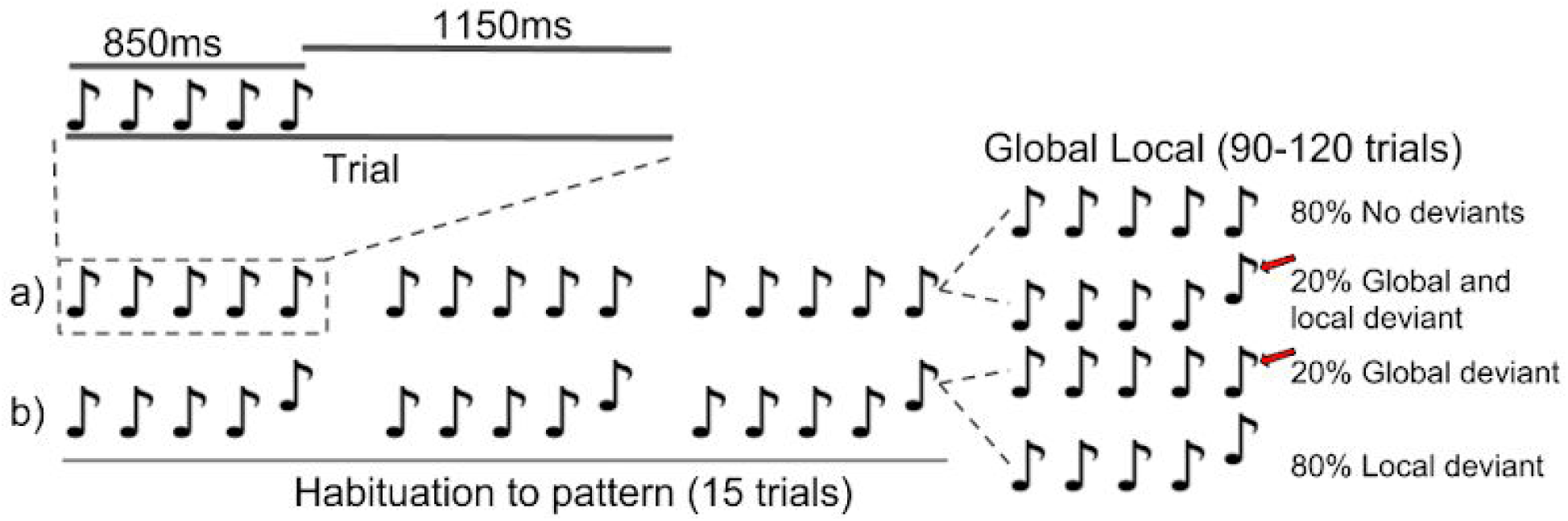
The “global-local” paradigm. One block consists of 105-135 trials, where each trial has five tones, with the fifth tone deviating or not from the previous four. The first 15 trials of a block were habituation trials and followed the same pattern; (a) no deviant fifth tone, or, (b) deviant fifth tone. The next 90-120 trials had a 20% chance to deviate from the pattern established in the habituation part of the block (red arrow).

#### Working memory task

In order to induce a high cognitive load, we designed a working memory task where participants were instructed to memorize then maintain a string of letters (consonants) before indicating which of three strings of three consonants were present in the original string. The sequence of answers and letters was randomized to minimize sequencing strategies. A trial consisted of a pause screen (until button press), a 3 second countdown, presentation of the text string for 10 seconds (update phase), fixation cross for 20 seconds (maintenance phase), and finally followed by three answer alternatives. The answer screen was present until answer was given. In addition, to ensure high subjective cognitive load we employed a staircase procedure so that each correct answer increased query string by one consonant (up to a total of 20), while a wrong answer decreased by one (down to a total of 3). The text was white on a uniform grey background. See Figure 2. The paradigm lasted for as long as necessitated by the measures.

**Figure 2.**
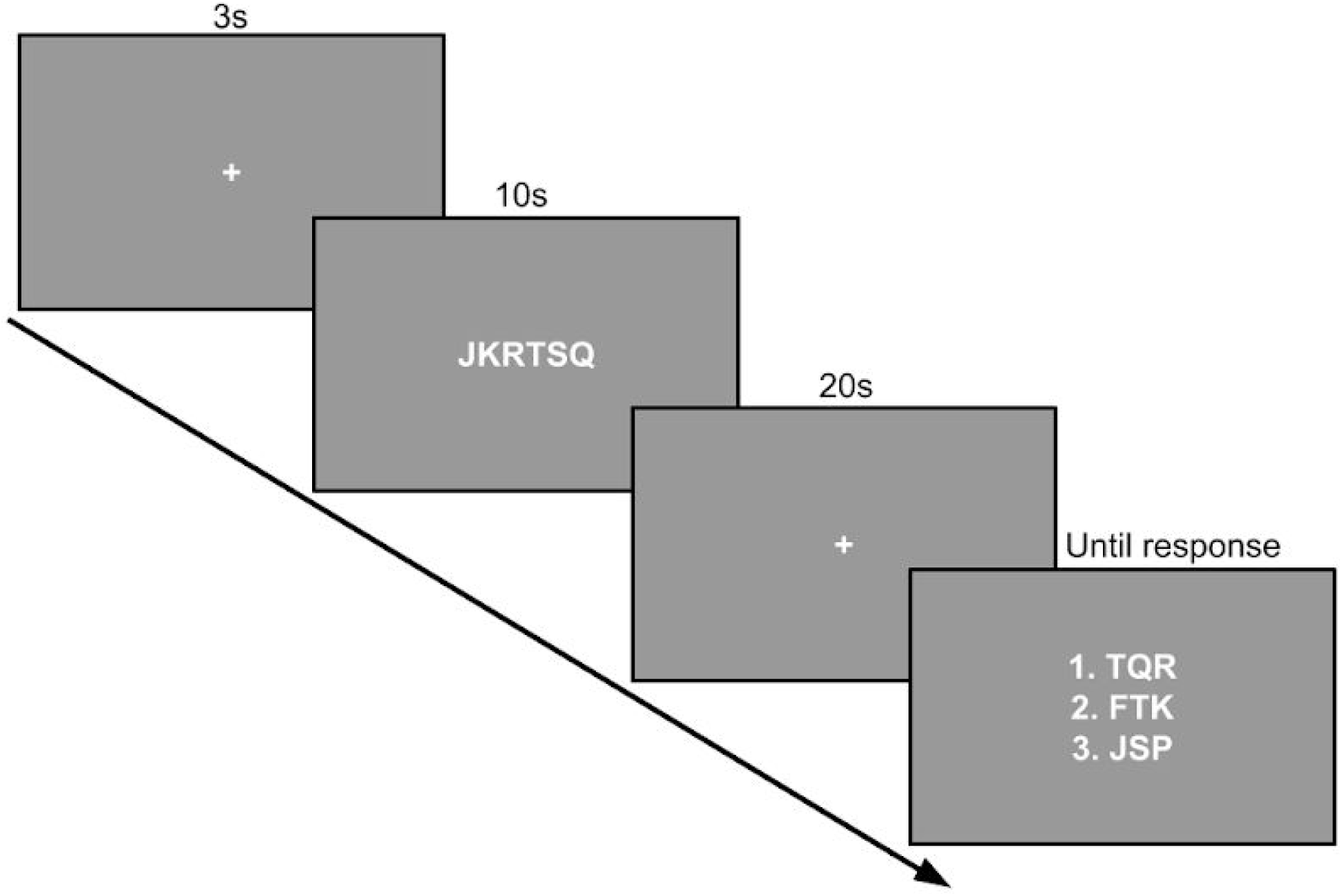
Working memory paradigm employed for attentional and cognitive load. One trial consisted of a fixation cross, a string of consonants (update phase), fixation cross (maintenance phase), then three response alternatives (response phase). The answer options consisted of three consonants each, were the two wrong alternatives contained one wrong letter (not present in the original string).

#### Low-load attentive task

For each of the three methods (TMS-evoked, auditory-evoked, and spontaneous EEG) we employed a low-load attentional paradigm consisting of merely paying attention to the stimuli. In the global-local auditory paradigm (for evoking the P3b), participants were asked to count the number of global deviants they heard, and report the number after each block of stimuli. For the TMS we asked participants to count the number of TMS pulses in total, and for the spontaneous EEG we asked participants to count the number of seconds (audible clicks from an analog wall mounted clock).

### Equipment/Setup

The anatomical MRI recorded for TMS-EEG navigation purposes was scanned at Philips Ingenia 3T scanner (Intervention Center, University Hospital Oslo). The sequence employed was a T1 weighted sequence with 1.0 mm * 1.1 mm voxels with 184 slices and 2.0 mm thickness, reconstructed to 256*256*184 matrix, SENSE=1.5 (RL), TR=4.6 ms, TE=2.3 ms, for a total sequence length of ∼4 minutes. The sequence was preceded by a B1 calibration scan.

For TMS stimulation we used 70mm cooled coil (PMD70-pCool) together with PowerMAG Research 100 stimulator (Mag&More, 81379 Munich, Germany). For 3D navigation, we used NDI Polaris Spectra spatial navigation system employing two infrared cameras, motion trackers, and a 3D reconstruction of participants head and brain (PowerMag view v1.7.4.401).

Resting motor threshold (RMT) was estimated using a Mobi MINI TMSi Bluetooth connected amplifier in conjunction with PowerMag control software. Electrodes for the RMT device were fixed in order to record muscle activity from the abductor pollicis brevis muscle on the dominant hand, in response to single TMS pulses to the thumb area of the contralateral motor cortex. RMT was estimated using an automatic algorithm that calculates the stimulation intensity that has the maximum likelihood to induce 50mv peak to peak electromyographic (EMG) response to single TMS pulses in 50% of the stimulations (Mishory et al., 2004).

The EEG was acquired with two BrainAmp DC 32 channel amplifiers and a 64 channel TMS compatible passive electrode cap (Brainproducts, 64Ch-EasyCap for BrainAmp with ECI electrode gel and NuPrep abrasive skin-prep gel). Acquisition was done with 5kHz sampling rate, DC-1000Hz hardware filter, 16 bit rate, and ±16.384mV measurement range.

The working memory and global local paradigms were programmed in Python 2.7.12 using PsychoPy2 (v1.82.02). The paradigms were run on a Dell latitude 3550 laptop running Xubuntu 16.04. The working memory paradigm was presented on a screen approximately 140cm in front of the participant’s head (FlexSan 2768, 19”, 60hz, 1024×768, Eizo Nanao Corp.), with participants using a keyboard in their lap with keypad buttons “1”,”2”,”3” for each response alternative.

The stimulation protocol used for TMS-EEG for the PCI was similar to that of Casali et al. (2013), with a series of 300 pulses (mean inter pulse interval 2 [±0.3s random jitter], full waveform) targeted at either the prefrontal cortex (BA6) or the parietal cortex (BA7) of the left hemisphere, with a stimulator intensity between 120-160% of the RMT. Targeting was aided with 3D spatial navigation employing motion trackers, and 3D reconstruction of the participant’s skull and brain based on the participant’s own MR-image. We adjusted intensity, location, and orientation of the induced magnetic field to maximize the peak-to-peak amplitude of the initial deflection of the TMS evoked potential (TEP, 20-40ms) and to minimize the level of artefacts present in the TEP. The goal was a minimum of 10mV peak-to-peak amplitude in the electrodes close to the stimulation area, and minimal artefacts in all electrodes.

To mask TMS noise during TMS-EEG recordings, we used continuous train of waveforms containing multiple TMS coil clicks of different intensity. TMS pulses was chopped into ∼1ms subsections and then randomly shuffled in time. This generated a noise-like sound capable of masking the audible click from the TMS pulse at quite low volume. During the TMS experiments, the volume of the masking sound was set high enough (within comfort range) to completely mask the sound generated by TMS pulses (perpendicular to cortical stimulation). The auditory stimuli for the global local paradigm consisted of 50ms duration (7ms rise and fall time) chords of 3 sinusoids (either 350, 700, and 1400 Hz; or 500, 1000, and 2000 Hz - second and third partials were of 1/2 and 1/4 intensity, respectively). During the global-local experiments, the volume of the sounds was adjusted to what participants considered clearly audible speech level. Auditory stimulus was run on a Dell latitude 3550 (Windows 8) using Sennheiser earplugs.

All preprocessing and analysis was performed using Matlab (M2017b) employing in house scripts, and EEGLAB (v14.1.1).

### Protocol

We first performed a standardized MRI for spatial navigation in TMS-EEG. We then investigated participants’ approximate RMT and degree of muscle activation in response to TMS pulses (in BA6, BA7). Only participants with RMT up to 70% of stimulator capacity and little to no muscle activations were included in the main study.

In the main study, first the EEG cap was mounted and electrodes adjusted with ideal target impedances of < 5kOhm. Following setup, participants were seated in a reclining chair, and then received 3 training trials of the working memory paradigm to minimize initial training effects and establish starting difficulty. We then performed the auditory paradigm (P3b), spontaneous EEG paradigm (LZc, ACE, SCE, DTF), and the TMS-EEG paradigm (PCI), counterbalanced between participants. For each method, participants first performed the working memory task (distracted high-load condition) followed by the passive attentive counting task (attentive low-load condition) to avoid anchoring effects. See Figure 3.

**Figure 3.**
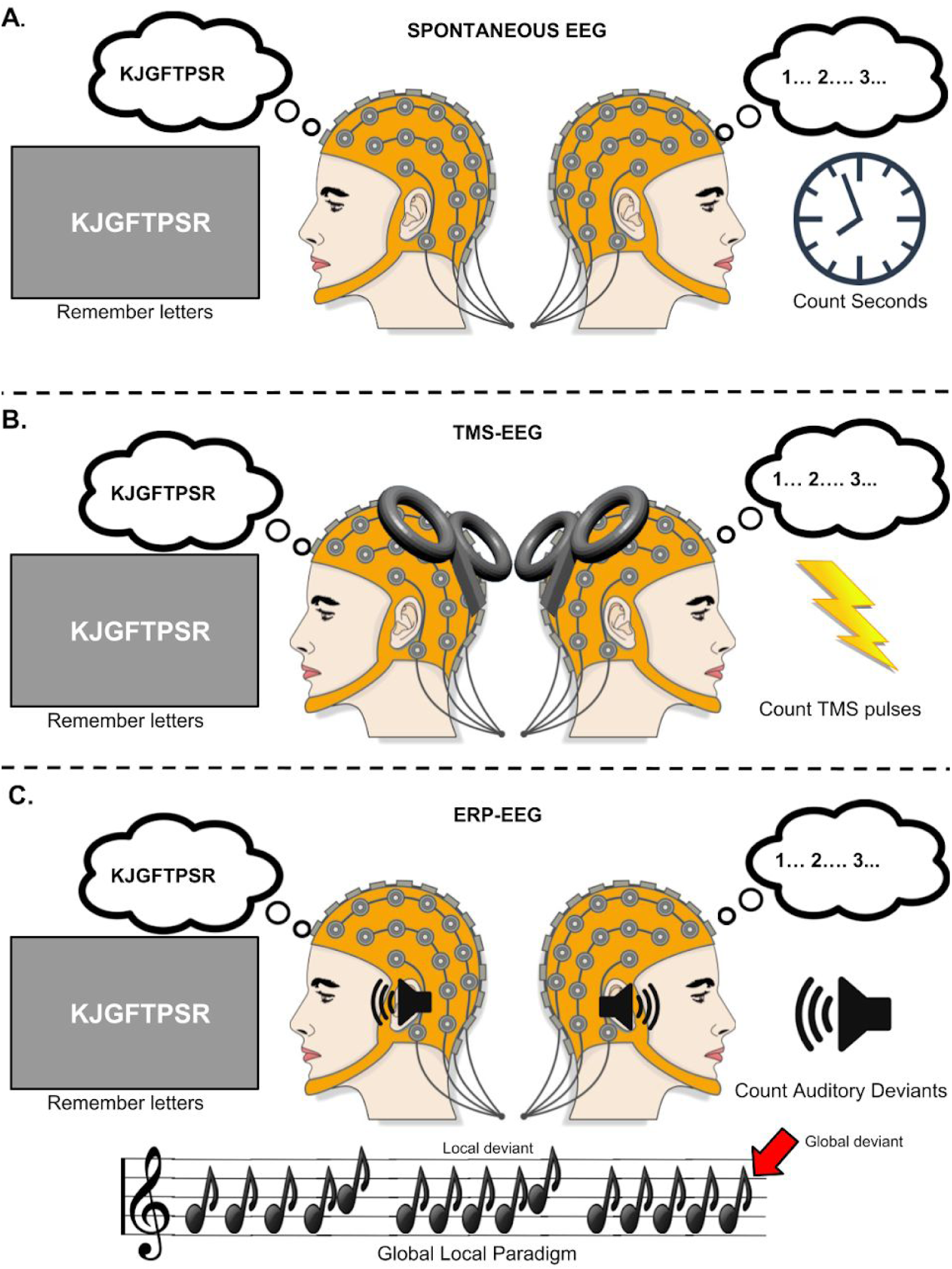
Overview of experimental paradigm. Three pseudo-randomized blocks with two conditions (low-load stimuli attentive, high-load working memory). **(A)** Spontaneous EEG recording while counting seconds, and while performing working memory task; **(B)** TMS-EEG recording while counting TMS pulses, and while performing working memory task; **(C)** ERP-EEG recording while counting number of global auditory stimuli pattern deviations in the global local paradigm, and while performing working memory task.

To control for attention in the auditory paradigm, we asked participants after both high and low-load conditions to indicate which of 10 specific auditory sequences they had heard and how certain they were (see Supplementary Materials S1).

For the spontaneous EEG recordings, participants did the high-load condition for 10 trials, then estimated how many seconds they believed had passed. Afterwards they performed the low-load condition, counting seconds. Participants were instructed to report how many seconds had elapsed afterwards (∼5-6 minutes).

For the TMS method, we first performed an RMT estimation to guide initial stimulator intensity. After finding a suitable location for stimulation, coil rotation, and stimulation intensity, the participant performed the high-load condition with 5 TMS pulses in the update phase, and 10 pulses in the maintaining phase of the paradigm. After 20 trials (300 pulses), we asked the participants to estimate how many pulses they had received. In the passive condition we delivered a continuous train of 300 pulses lasting for a total of 10 minutes which participants were instructed to count.

### Preprocessing

Preprocessing followed similar steps as in previous studies (DTF; Juel, Bremnes, et al., 2018. PCI; Casali et al., 2013. ACE, SCE, LZc; Schartner et al., 2015) with some deviations. See Figure 4 for details. Novel to this study, for the PCI measure, we first removed the pulse artefact (−1 to 10ms around each TMS trigger), and replaced it with normally distributed white noise with mean and standard deviation of the baseline (−300 to −50ms). For the spontaneous measures, we created non overlapping windows of 5000ms length, and limited the number of electrode channels used in the analysis (after preprocessing) to 9 (F5, Fz, F6, C5, Cz, C6, P5, Pz, P6) to ensure that each 5 seconds window (3000 samples after downsampling) would have enough samples to estimate a probability distribution of 2^channels=512 possible states. The channels were selected based on broad topological coverage.

In general for all methods, we implemented an automatic trial cleaning method where epochs were individually rejected (20% highest scoring within condition) using an automatic classifier based on a “noise” scoring algorithm:

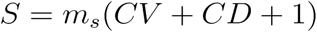

Where *S*=score, *CV*=number of channels with over 200uV peak-to-peak amplitude within the epoch, *CD*=number of channels with instantaneous amplitude change above 100uV at least once in the epoch, and *m*_*s*_=mean of channel variances. This equation scores high with high variance, modulated by the number of channels showing unrealistic amplitudes or sharp deflections. The results of the automatic scanning algorithm were independently validated by two researchers performing manual classification on two randomly chosen subjects.

### Analysis

#### P3b

The epoched data from the global-local paradigm were analyzed following Bekinschtein et al. (2009). Here we focus on the channel “Pz” as it’s considered the dominant channel for the global effect (King et al., 2013) in the time window 300-700ms after the onset of the fifth sound. First, we averaged across trials according to the four trial types; Local Standard (LS), Local Deviant (LD), Global Standard, (GS) Global Deviant (GD). As we were only interested in the global effect (GD vs GS), the global effect was calculated as a t-statistic at each timepoint in the ERP of GD vs GS. This resulted in an ERP of p-values for each channel. The p-values were set to “0” if lower than the peak (lowest) p-value in baseline (−1000 to −200ms), and if *p*<.01 for at least 10 consecutive timepoints (25ms). Discarded p-values were replaced with “1”. These binary values were then averaged. This resulted in a single value for each subject, where a value of “1” indicated that there was no significant global effect according to the above criteria, and a value of < 1 indicated some significant global effect.

#### PCI

For calculating PCI we calculated the inverse solution using minimum norm estimate based on the MNI ICBM152 standard atlas MRI. Then the statistically significant source activity was estimated using a bootstrap resampling of the baseline activity. Briefly, a source was given a value ‘1’ at some point in time if its activity at that time was larger than the 99th percentile of the maximal amplitudes in the bootstrap resampled baselines. Otherwise, the source was given a value ‘0’. This yielded a 2D binary matrix representing the spatiotemporal activity patterns of the cortical sources in response to a TMS pulse, from which LZc was calculated on the interval 15-300ms, concatenated in the spatial dimension. The complexity was then normalized based on the analytic asymptotic peak LZc for a signal of a given length and source entropy to yield the PCI. For details see supplementary materials of (Casali et al., 2013).

#### Spontaneous EEG measures

The DTF was calculated using the ‘DTF’ function in the eConnectome toolbox (He et al., 2011), and followed the Juel et al (2018) with some deviations. Specifically, DTF was calculated using 600 Hz sampling rate, 9 channels, and 5 second epochs (instead of 512 Hz, 25 channels, and 1 second epochs). The deviations has been shown in unpublished analysis to minimally impact results while reducing computational demands and relying on similar preprocessing as the other spontaneous measures. To produce a one-dimensional metric, we took the median of the logarithm of DTF values over epochs and frequencies for each channel pair, resulting in a matrix of median log(DTF) connectivity for all channel pairs. This matrix was further compressed to obtain the median information outflow from each channel. Based on this, the root mean square difference between all values of median information outflow was calculated for each subject, yielding a single value termed DTF heterogeneity (DTF_HET_), indicating the heterogeneity of ‘information outflow’ across channels for a given subject within a condition.

For the ACE and SCE we first performed a Hilbert transformation, then binarized the data. For ACE, we used a median split of the amplitudes, setting any time point with amplitude above the median of that channel in the epoch to ‘1’ (‘0’ otherwise). For SCE, the threshold was based on the synchrony between each pair of channels, so that a channel pair was considered to be in synchrony and given a value ‘1’ if the difference between their instantaneous phases was below 0.8 radians (‘0’ otherwise). The resulting binary matrices represented amplitude of activity for the set of channels over time (ACE), or the states of phase synchrony between channel pairs over time (SCE). Based on these matrices, we found the distribution of states over time where a state (or ‘coalition’) was defined as a the pattern of 1’s and 0’s across channels at one time point. From the state distribution we then calculated the sample entropy and normalized with respect to the mas entropy of 50 shuffles of the original data (Schartner et al., 2015). Our final measure for the ACE was the mean ACE values over epochs (for each participant in each condition), while the final SCE value was the mean SCE over channels and epochs.

The LZc followed the same steps for binarization as for the ACE. However, here, the complexity was quantified by calculating the LZc (using the algorithm presented in (Kaspar & Schuster, 1987)). Each LZc value was normalized with respect to the complexity of the shuffled original data (maximum of 50 shuffles). The final value for the LZc for a participant in a condition was the mean LZc values across epochs.

We then performed the same statistical analysis for all measures. First, we calculated the rate of true positives and false negatives according to previously calculated thresholds for each measure (see source articles: Bekinschtein et al., 2009; Casali et al., 2013; Juel, Bremnes, et al., 2018; Schartner et al., 2015). Since participants were considered to be conscious in both of our conditions, the ground truth for all the participants state was always positive. Thus, classifications were either false negatives or true positives. Secondly, we performed pairwise and two-sample t-tests to test whether there was any significant difference between the low and high-load conditions. Third, to account for potentially non-normal distributions of the measures in each condition, we performed a permutation (2^12^ = 4096 permutations) test with condition as the independent variable (see Karniski, Blair, & Snider, 1994). Fourth, we implemented a simple data driven classification algorithm to test if we could correctly classify the participants as being attentive (low-load) or distracted (high-load). For the classifier, we first constructed distributions for each condition (for all participants minus one). Then, we calculated the probability that the values observed for the given participant came from the attentive distribution:

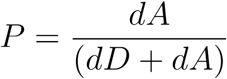

where *P* is probability of datapoint coming from the attentive condition, *dA* is standardized distance from low-load condition distribution, *dD* standardized distance from high-load condition distribution. This results in 0<*P*<1 where *P*>.5 indicates that the datapoint is more likely to come from the low-load condition, and *P*<.5 indicates the opposite. Fifth, and finally, we calculated an ROC curve based on thresholding the probability distributions observed in step 4 to investigate whether a cut off could be found that separated the two conditions. Step 4 and 5 were performed to test for single subject classification.

## Results

The effects of attentional and cognitive load can be seen in Figure 5 and Table 1. Specifically, we observed that in the low-load (i.e. attentive) condition (marked by blue triangles, 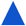, in Figure 5a,b), all measures were above pre-established thresholds (red dashed lines, - -, in Figure 5A, **θ** in Table 1) that have been used to separate conscious vs. unconscious states in previous studies (see Casarotto et al., 2016; Juel, Bremnes, et al., 2018; Schartner et al., 2015), except some participants who scored below threshold for ACE (*n* = 2) and DTF_HET_(*n* = 2). For the high-load (i.e. distracted) condition (marked by green circles, 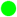 in Figure 5a,b), all participants obtained LZc, ACE, SCE, PCI and DTF_HET_ scores above the pre-established thresholds, except for some participants who scored below threshold for DTF_HET_ (*n* = 2). In contrast, all scores of the P3b global effect fell below the previously established threshold (King et al., 2013), as shown in Figure 5A, right panel. As participants were conscious in both the low and high-load conditions, scores above pre-established thresholds were considered true positives, while scores below thresholds were considered false negatives. For DTF_HET_ both participants who scored below threshold in the low-load condition were above the 95% confidence interval for the anesthetic state (Xenon and Propofol), while one of the two participants scoring below threshold in the high-load condition were above the 95% confidence interval (Juel, Bremnes, et al., 2018). For ACE, both participants who scored below pre-established threshold were above the 95% confidence interval for anesthetic state (Propofol; Schartner et al., 2015). This suggests that all the below threshold values in DTF_HET_ and ACE (except one) can be considered borderline cases, and that pre-established thresholds need a broader norm base than currently offered.

**Table 1.** Measures and their respective preprocessing steps. Removal of bad channels consisted of manual inspection, with a focus on dead channels or channels with consistent high amplitude artefacts relative to the other channels. Removed channels were interpolated. Independent Component Analysis (ICA) was used to decompose the signal into independent components (ICs), which were manually inspected and removed if they appeared to be associated with ocular or pulse related artefacts. Epoching was centered on the TMS pulse, the fifth auditory stimuli within a series (see analysis), or continuous non-overlapping windows, depending on method. Baseline subtraction consisted of removing the average amplitude of the specified baseline relative to the epoch center/zero point.

| MEASURES |  |  |
| --- | --- | --- |
| P3B | PCI | DTF, ACE, SCE, LZc |
| METHOD |  |  |
| AUDITORY | TMS | SPONTANEOUS |
| PREPROCESSING |  |  |
| 1. Remove and interpolate bad channels<br>2. Downsampling 250Hz<br>3. Bandpass filter 0.5Hz to 100Hz<br>4. Average reference<br>5. ICA decomposition<br>6. Ocular artefact ICs removed<br>7. Lowpass filter 20hz<br>8. Epochs $\pm 1000$ms around fifth sound<br>9. Baseline subtraction [-1000, -50]ms<br>10. Epoch rejection | 1. Remove & replace pulse artefact [-1,10]ms<br>2. Remove and interpolate bad channels<br>3. Downsampling 362.5Hz<br>4. Bandpass filter 0.1Hz to 45Hz<br>5. Average reference<br>6. ICA decomposition<br>7. Ocular artefact ICs removed<br>8. Epochs [-300, 600]ms around TMS pulse<br>9. Baseline subtraction [-300,-50]ms<br>10. Epoch rejection<br>11. ICA decomposition<br>12. Pulse related ICs removed | 1. Remove and interpolate bad channels<br>2. Downsampling 600Hz<br>3. Bandpass filter 0.5Hz to 100Hz<br>4. Average reference<br>5. ICA decomposition<br>6. Ocular artefact ICs removed<br>7. Lowpass filter 45hz<br>8. Epochs $\pm 2500$ms, non-overlapping<br>9. Baseline subtraction [$\pm 2500$ms]<br>10. Surface Laplacian filter*<br>11. Channel selection |

\*Only for DTF

**Table 2.**
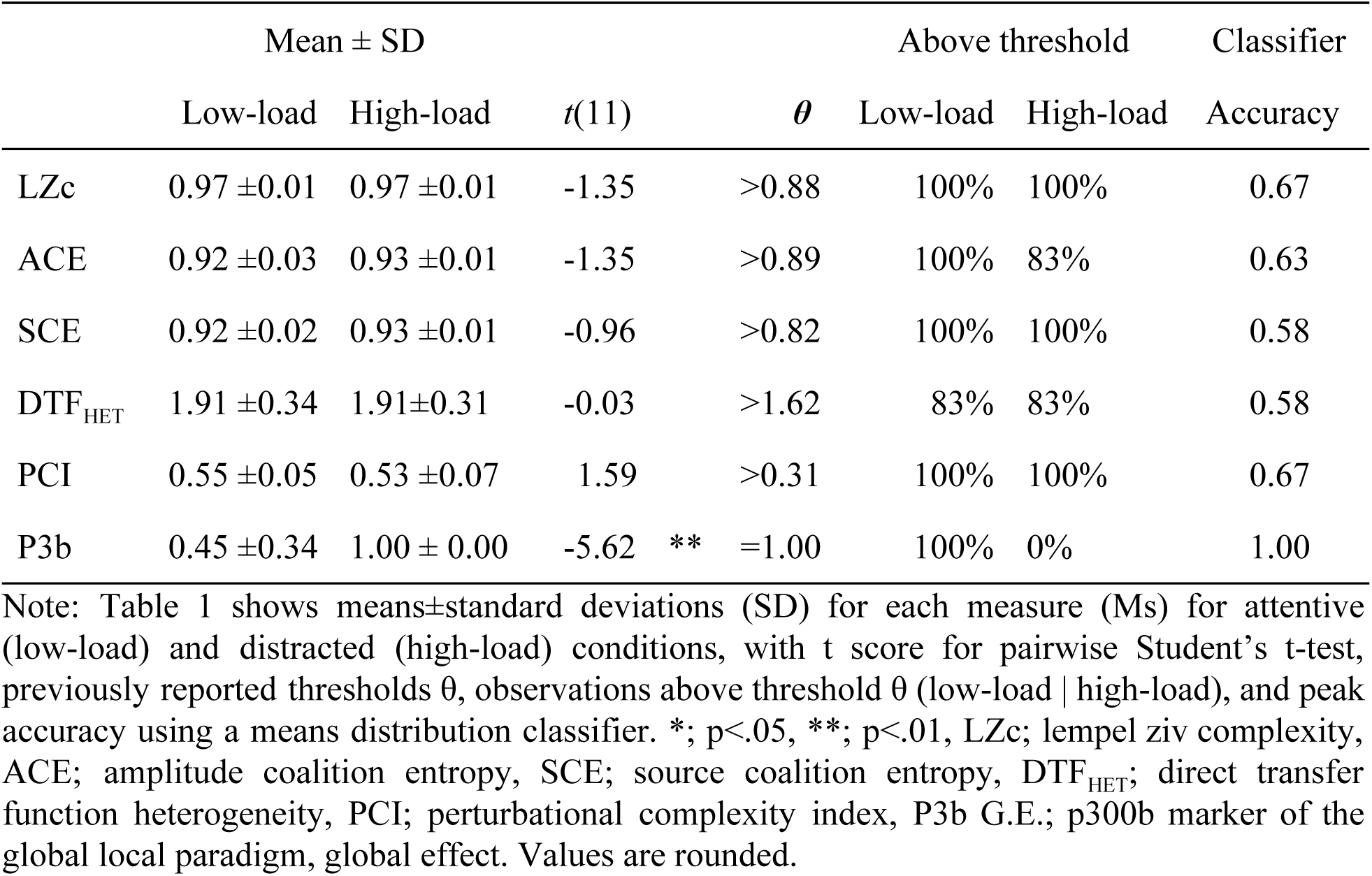
Descriptive statistics and results for each measure.

| | Mean ± SD | | | $\theta$ | Above threshold | | Classifier |
| --- | --- | --- | --- | --- | --- | --- | --- |
| | Low-load | High-load | $t(11)$ | | Low-load | High-load | Accuracy |
| LZc | 0.97 ± 0.01 | 0.97 ± 0.01 | -1.35 | >0.88 | 100% | 100% | 0.67 |
| ACE | 0.92 ± 0.03 | 0.93 ± 0.01 | -1.35 | >0.89 | 100% | 83% | 0.63 |
| SCE | 0.92 ± 0.02 | 0.93 ± 0.01 | -0.96 | >0.82 | 100% | 100% | 0.58 |
| DTF<sub>HET</sub> | 1.91 ± 0.34 | 1.91 ± 0.31 | -0.03 | >1.62 | 83% | 83% | 0.58 |
| PCI | 0.55 ± 0.05 | 0.53 ± 0.07 | 1.59 | >0.31 | 100% | 100% | 0.67 |
| P3b | 0.45 ± 0.34 | 1.00 ± 0.00 | -5.62 ** | =1.00 | 100% | 0% | 1.00 |

Note: Table 1 shows means ± standard deviations (SD) for each measure (Ms) for attentive (low-load) and distracted (high-load) conditions, with $t$ score for pairwise Student's $t$ -test, previously reported thresholds $\theta$ , observations above threshold $\theta$ (low-load | high-load), and peak accuracy using a means distribution classifier. \*, $p < .05$ , \*\*, $p < .01$ , LZc; lempel ziv complexity, ACE; amplitude coalition entropy, SCE; source coalition entropy, DTF<sub>HET</sub>; direct transfer function heterogeneity, PCI; perturbational complexity index, P3b G.E.; p300b marker of the global local paradigm, global effect. Values are rounded.

**Figure 5.**
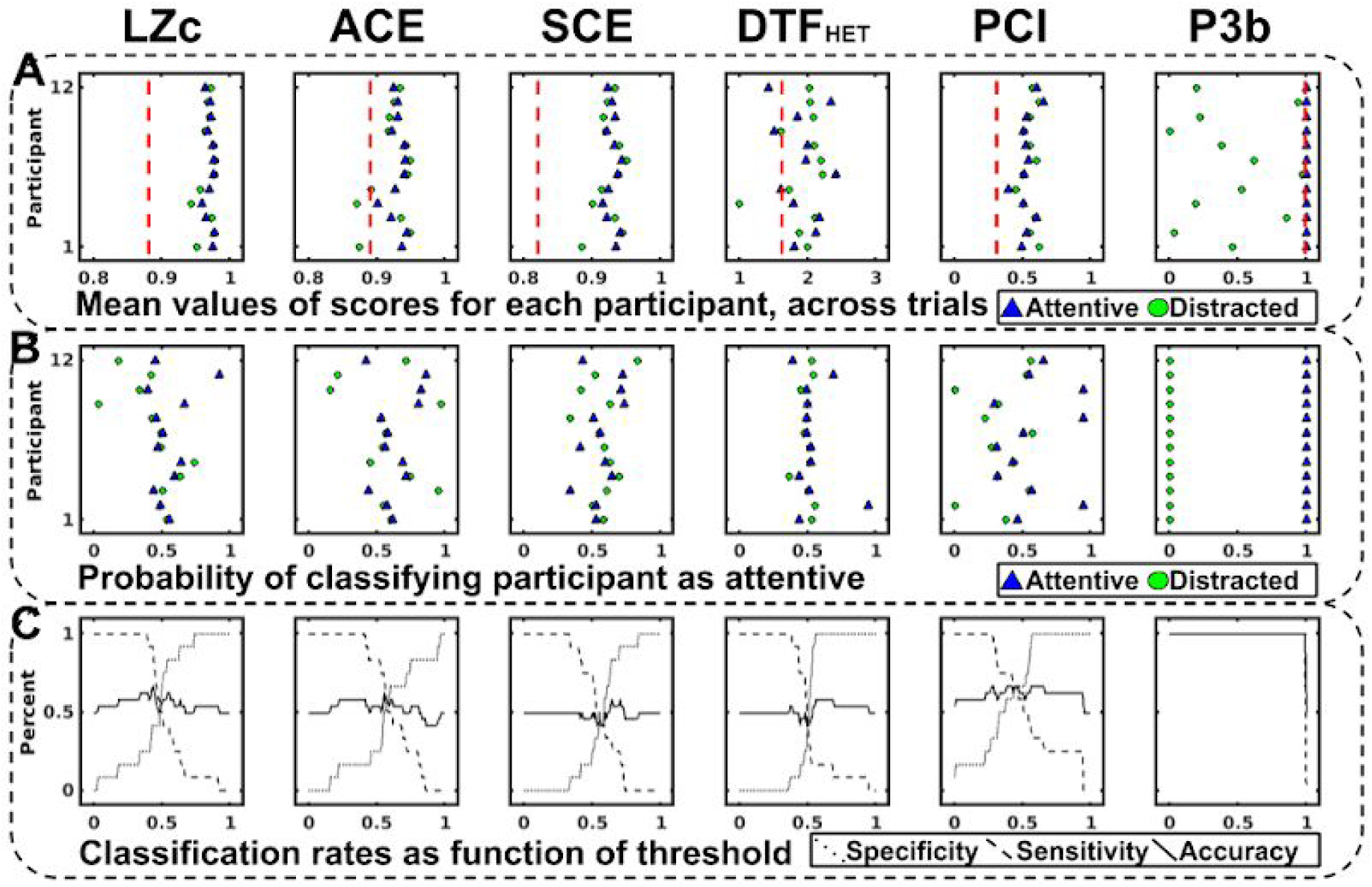
Overview of results. Results for each measure, ACE, PCI, DTF_HET_, SCE, LZc, and P3b, for the distracted (high-load) and attentive (low-load) conditions. **(A)** Mean score for each measure under each condition for each participants. The red dashed lines in 5A indicate the pre-established thresholds that have previously been used to indicate consciousness **(B)** The probability of being classified as “attentive condition” given the overall distribution of the group (equation 2) for each participant in each condition. **(C)** Sensitivity, specificity, and accuracy of varying thresholds θ, based on B.

Finally, we observed no significant statistical difference (following pairwise and permutational statistical tests) between the low-load and high-load conditions for ACE, SCE, LZc, PCI, and DTF_HET_. However we did observe significantly higher P3b values for the high-load condition (*M*=1.00 ±0.00) than the low-load condition (*M*=0.45 ±0.34), two-samples *t*(11)=5.62, *p*<.01. P3b was also the only measure where the two conditions could be reliably separated on an individual basis by a classifier (Figure 5b) and by means of a threshold (Figure 5c). In sum, this indicates that only the P3b measure can reliably separate a low-load (attentive) and high-load (distracted) condition on both a group and single subject level, and is as such affected by loading of attentional and cognitive resources by a working memory task in lieu of attending to the auditory global-local paradigm itself.

## Discussion

In the present study we tested whether attentional and cognitive load affected some recently proposed EEG-based measures of state of consciousness in humans. In summary, our results showed that the ACE, SCE, LZc, PCI, and DTF_HET_ were not measurably affected by attentional or cognitive loading, suggesting that previous results are not substantially confounded by such effects (Casali et al., 2013; Juel, Romundstad, et al., 2018; Schartner et al., 2015). In contrast, the P3b showed a 100% ability to discriminate between the two conditions, an effect which is even more pronounced in our data compared to previous findings from studies in which attention was modulated (Bekinschtein et al., 2009; King et al., 2013). This might indicate that the working memory task is a stronger cognitive loading task than those previously employed. Regarding the supplementary measures, they indicated that the participants were indeed focused on the working memory paradigm and did not pay much attention to stimuli or durations (see Supplementary S1 and S2).

The present study was primarily designed as a quality control study of the measures tested, as any significant difference between the effects of distracting cognitive load and passive attention to stimuli or duration would make the proposed measures weaker in terms of classifying consciousness as such.

Of secondary interest, we note that several of the proposed measures (ACE, SCE, LZc, and DTF_HET_) could potentially be employed as real-time clinical measures or monitors of the state of consciousness in humans due to their high temporal resolution and continuous output over an extended period of time. This may allow for automatic bedside processing and classification.

However, there are some limitations regarding the results and interpretations. First, it could be argued that the low-load counting tasks employed did not only differ in degree of cognitive load but also in sensory processing and modality (counting vs. working memory). To test for similarity in tasks, we re-analyzed data from a passive mind wandering condition (N=6) from a different experiment employing the same technical setup (Farnes, Juel, Nilsen, Romundstad, & Storm, 2019), and observed similar values as presented in this paper, within ∼1 SD of the distribution of the attentive, low-load condition (*M*_*ACE*_=0.898, *M*_*SCE*_=0.91, *M*_*LZ*_=0.91). In addition, a PSD analysis replicated previously observed increases in theta band power in frontal electrodes (channel Fz in particular) during working memory modulation (Hsieh et al., 2011; Hsieh & Ranganath, 2014), indicating that the high-load condition did indeed modulate working memory (see Supplementary materials S4). While this doesn’t exclude that the low-load attentive and high-load distracted task differed in more ways than cognitive load, the results suggests that the overall goal of investigating within state (awake) cognitive load modulations remains valid.

Secondly, for ACE and DTF_HET_ we observed some (*n*=2/12 and 4/12) results below the pre-established threshold values of conscious vs. unconscious states employed previously (Juel, Bremnes, et al., 2018; Schartner et al., 2015). However, while the subjects were clearly conscious, these values were relatively close to the thresholds compared to typical values observed during unconsciousness, all observations (except one for DTF_HET_) were above respective 95% confidence intervals for the anesthetic state. However, these low values relative to the threshold, might be related to three broad possible limitations; (i) difference in acquisition and preprocessing parameters from previous studies, (ii) selection of channels used for the final analysis, and (iii) the basis for threshold estimation. For the first factor (i), it can be argued that since there exists 2^channels^ number of available states (coalitions) one needs to sample as much of the full probability distribution as possible. This places limits on number of channels and samples as one needs enough samples in one epoch to have high probability to sample the maximum entropy probability distribution of states/coalitions (see Supplementary materials S3). While Schartner et al. (2017) dismissed this issue, we could not find any strong theoretical or empirical reason to do so. As such, we limited analysis to 9 channels and 5 second epochs with 600Hz sampling rate. For the second factor (ii), it’s conceivable that the subset of channels chosen affects the absolute values of the measures. While more rigorous investigation of complexity as a function of channel selection is warranted, we focused on broad topological coverage instead of a specific subregion of cortical channels as there’s no strong arguments for favouring a local area as a basis for calculations. For the third factor (iii), it might be the case that demographic factors, differences in technical and practical implementation, or deviations in the analysis and data cleaning could warrant that a new cutoff be calculated based on a sample gathered locally. Thus, the exact results from these three methods may depend on details of preprocessing, equipment, and other confounding variables.

Finally, while the present study did not employ a non-conscious control, it’s important to perform studies focusing on validating measures within conscious states to control for possible confounding factors such as muscle tension and memory. Muscle tension in particular has been shown to influence metrics of anesthetic depth such as the bispectral index (Schuller, Newell, Strickland, & Barry, 2015), and loss of consciousness during anesthesia could be confounded by partial amnesia, since it depends on self-reports after waking up, and several general anesthetics are known to cause partial amnesia. For a discussion of anesthesia and unconsciousness, see (Sanders, Tononi, Laureys, & Sleigh, 2012). To control for such effects, measures of conscious state should be investigated in for example patients with amyotrophic lateral sclerosis (ALS) as they have non-abnormal muscle tension (Rowland, 1998), and patients with anterograde amnesia as in Korsakoff’s syndrome (Weiner & Craighead, 2010). Future studies should also expand beyond merely the conscious and unconscious divide, but additionally investigate measures in relation to memory formation, muscle tone, altered states of consciousness, and clinical populations with severely altered phenomenological consciousness. One can in this way arrive at a set of candidate measures that are validated over a broad range of conditions, including modulations within conscious and unconscious states. Here we showed that the candidate measures PCI, ACE, SCE, and DTF_HET_, were not affected by attentional and cognitive loading, and are thus unlikely to depend on subject task compliance.

## Funding

This study was supported by the European Union’s Horizon 2020 research and innovation program under grant agreement 7202070 (Human Brain Project (HBP) SGA1 grant and the Norwegian Research Council (NRC: 262950/F20 and 214079/F20).

## Acknowledgments

We would like to thank PhD. Benjamin Thürer for critical feedback and corrections, Prof. Marcello Massimini for helpful suggestions and discussions, and the subjects for volunteering to participate in this study. We also would like to thank the Intervention Center at the Oslo University Hospital for MRI and lab facilities.

## Author contributions and notes

A.S.N, B.E.J. and J.F.S. planned and designed the study. A.S.N. and B.E.J. performed the experiments and collected the data. A.S.N. and B.E.J. analyzed the data. A.S.N created figures and drafted the paper. A.S.N, B.E.J. and J.F.S. critically reviewed the paper. The authors declare no conflict of interest.

## Supplementary Materials

### S1

To control for the possibility that participants did attend to the sounds in the global-local auditory paradigm (used to calculate the P3b measure) even though instructed to pay attention to the working memory task (high-load, distracted condition), we implemented a sound sequence questionnaire. After completion of the global local paradigm for both cognitive load conditions (low-load counting and high-load working memory), participants answered for each sound series whether they had heard it or not, and how certain they were. Series 1,2,4, 7, and 9, where present in the paradigm.

Overall, following the high-load distracted condition, participants answered on average 66.6% correct, with a subjective certainty of 77.0/100.0. Following the counting condition, participants answered on average 84.2% correct, with a subjective certainty of 92.9/100.0. There was a significant difference in accuracy (*M*_HIGH-LOAD_=0.66, *SD*=0.06, *M*_LOW-LOAD_=0.84, *SD*=0.05, *t*(11)=-3.09, *p*=.01), and confidence (*M*_HIGH-LOAD_=77.03, *SD*=3.04, *M*_LOW-LOAD_=92.93, *SD*=1.98, *t*(11)=-5.23, *p*<.001). See Figure S1 and table S1.

**Figure S1.**
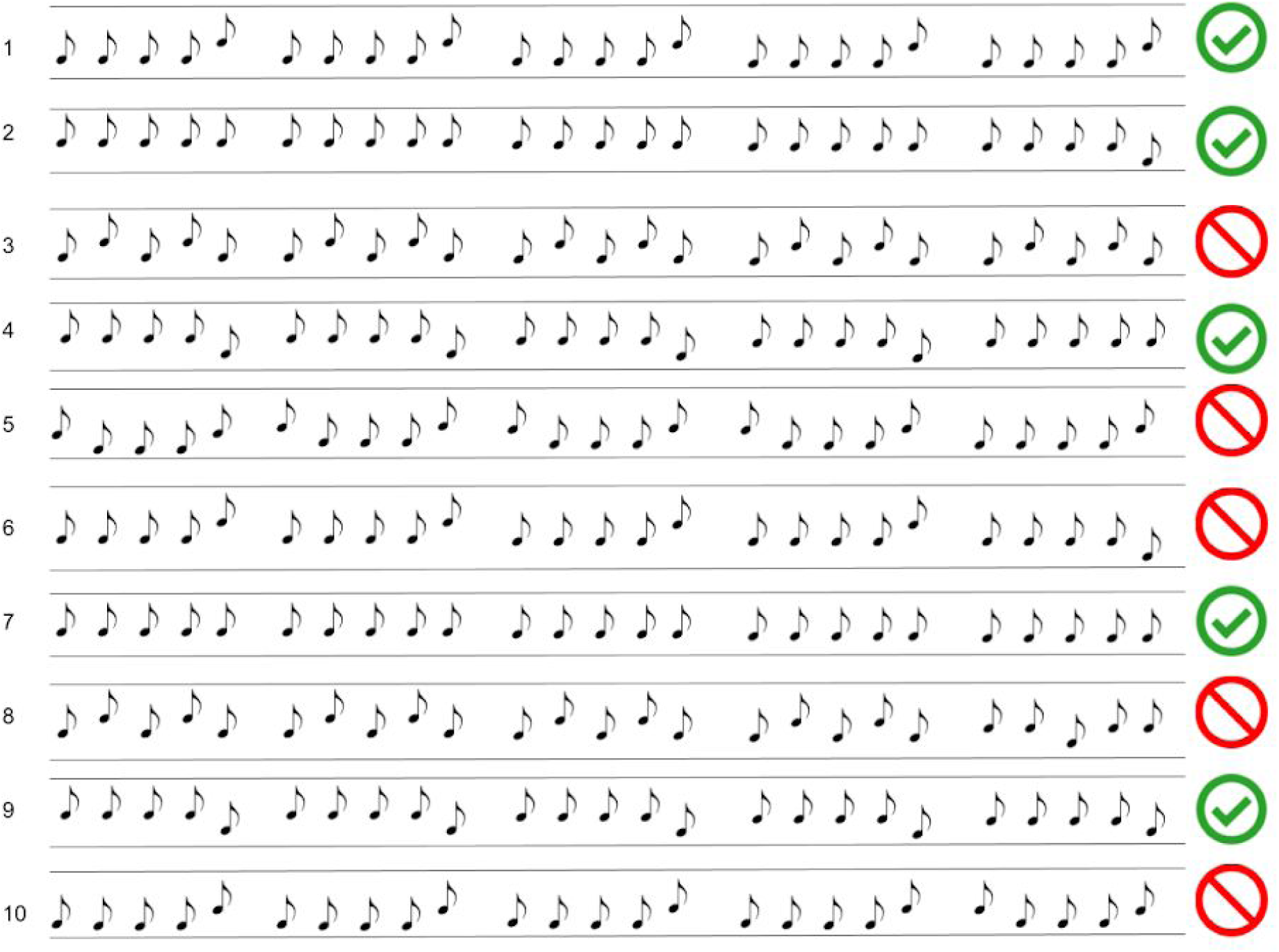
Sound Sequence Questionnaire.

### S2

In the counting condition, participants counted the number of TMS pulses, the number of seconds (using a wall mounted clock), and auditory global deviants in the global local paradigm. After the spontaneous EEG and TMS-EEG blocks, participants were asked to estimate how many seconds had passed and how many TMS pulses they had received, respectively. For the auditory paradigm, see supplementary materials S1. For the TMS-EEG measurements, participants estimated on average −137.25 pulses from target (SD=138.06) in the distracted high-load condition (2 participants within ±10% of target), and on average −1.75 pulses from target (SD=11.09) in the attentive low-load condition (12 participants within ±10% of target). Following the passive EEG paradigms, participants estimated on average −8.00 seconds from target (SD=259.17) in the distracted high-load condition (3 participants within ±10% of target) and on average 5.36 seconds from target (SD=9.22) in the attentive low-load condition (10 participants within ±10% of target). One participant failed to answer. See table S1.

**Table S1.** Overview of control parameters.

| Metric | Participant |  |  |  |  |  |  |  |  |  |  |  |
| --- | --- | --- | --- | --- | --- | --- | --- | --- | --- | --- | --- | --- |
|  | 1 | 2 | 3 | 4 | 5 | 6 | 7 | 8 | 9 | 10 | 11 | 12 |
| Accuracy Sounds, working memory (x/1) | 0,7 | 0,7 | 0,7 | 0,9 | 0,8 | 0,5 | 0,2 | 0,5 | 0,8 | 0,6 | 0,8 | 0,8 |
| Certainty Sounds, working memory (x/100) | 62,5 | 81,5 | 87 | 69 | 81 | 61 | 89 | 83 | 80 | 79 | 89 | 62,4 |
| Accuracy Sounds, counting (x/1) | 1 | 1 | 0,8 | 1 | 0,7 | 0,9 | 0,8 | 0,5 | 0,9 | 0,7 | 1 | 0,8 |
| Certainty Sounds (x/100) | 95 | 97 | 100 | 76 | 98 | 96 | 93,9 | 92 | 91 | 84,8 | 100 | 91,5 |
| Resting Motor Threshold (x/100) | 65 | 67 | 27 | 62 | 51.5 | 55.5 | 50 | 62 | 63 | 41.5 | 63,5 | 52 |
| TMS intensity (x/100) | 85 | 80 | 70 | 72 | 70 | 70 | ?? | 75 | 76.5 | 50 | 65 | 70 |
| TMS pulses estimation, working memory. | 500 | 200 | 300 | 70 | 40 | 300 | 50 | 50 | 80 | 20 | 150 | 200 |
| Target | 298 | 298 | 313 | 300 | 300 | 300 | 300 | 300 | 300 | 300 | 298 | 300 |
| TMS pulses estimation, counting. | 315 | 299 | 285 | 294 | 287 | 327 | 284 | 299 | 291 | 297 | 300 | 300 |
| Target | 306 | 299 | 300 | 297 | 298 | 300 | 300 | 300 | 300 | 300 | 299 | 300 |
| Seconds estimation, working memory. |  | 300 | 480 | 50 | 60 | 360 | 180 | 360 | 180 | 900 | 300 | 80 |
| Target |  | 377 | 483 | 291 | 245 | 335 | 285 | 205 | 175 | 197 | 239 | 206 |
| Seconds estimation, counting. |  | 348 | 200 | 272 | 250 | 180 | 243 | 220 | 176 | 200 | 270 | 203 |
| Target |  | 345 | 200 | 270 | 245 | 180 | 240 | 205 | 176 | 197 | 239 | 206 |

Note: Overview of TMS specific parameters and individual responses to estimations attached to each methodology and condition (high-load working memory, low-load counting). *Accuracy/certainty sounds* refers to Supplementary Figure S1; *Resting Motor Threshold* refers to the estimated required TMS intensity required to trigger a peripheral muscle activation 50% of the time, measured as x/100 where 100 is maximum TMS stimulator output; *TMS intensity* refers
to the stimulation intensity used in study; *TMS pulses estimation* refers to the subjectively estimated number of pulses received, with *target* referring to the actual number of pulses received; *Seconds estimation* refers to the estimated number of seconds elapsed during one session, with *target* referring to actual number of seconds elapsed.

### S3

To calculate the expected number of samples needed to fully sample a distribution at least once in each state (2^*n*^ states, with *n*=number of channels), one can use the inclusion-exclusion principle and De Morgan’s law, to solve the following to find the probability of getting at least one of each state given a set number of samples, *s*:

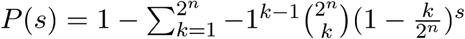

where *s*=number of samples to estimate probability of, *n*=number of channels/nodes. For 3000 samples (600Hz sampling rate and 5s epoc), we get the probability *P*(3000)=23.06%. Conversely, with 3000 samples one expects to sample ∼500 (of 512) possible states, and to get over *P*(s)>90% one needs ∼6000 samples. As such, while we’re on the lower end in this study, previous work has been far more liberal (Schartner, Carhart-Harris, et al., 2017; Schartner, Pigorini, et al., 2017; Schartner et al., 2015).

### S4

To validate that the working memory task had the intended effect we performed a power spectral density (PSD) analysis on the spontaneous EEG data using the Welch method (Welch, 1967), calculating PSD with a window of ⅔ of trial length (5s epoch) and ⅓ overlap, averaging across trials for each frequency bin, and finally calculating t-values for the pairwise difference of each frequency bin for each channel. According to previous literature (Hsieh & Ranganath, 2014), we expected an increase in theta power (4-7Hz) in frontal channels, dominantly channel Fz. We observed significantly higher mean power in the theta range (4-7 Hz) of channel Fz for the distracted condition relative to the attentive condition (*M*_DISTRACTED_=-15.72, *SD*=0.93, *M*_ATTENTIVE_=-16.48, *SD*=0.63, *t*(11)=2.90 *p*=.0298), indicating an effect of attentional loading, see Figure S2 and S3.

**Figure S2.**
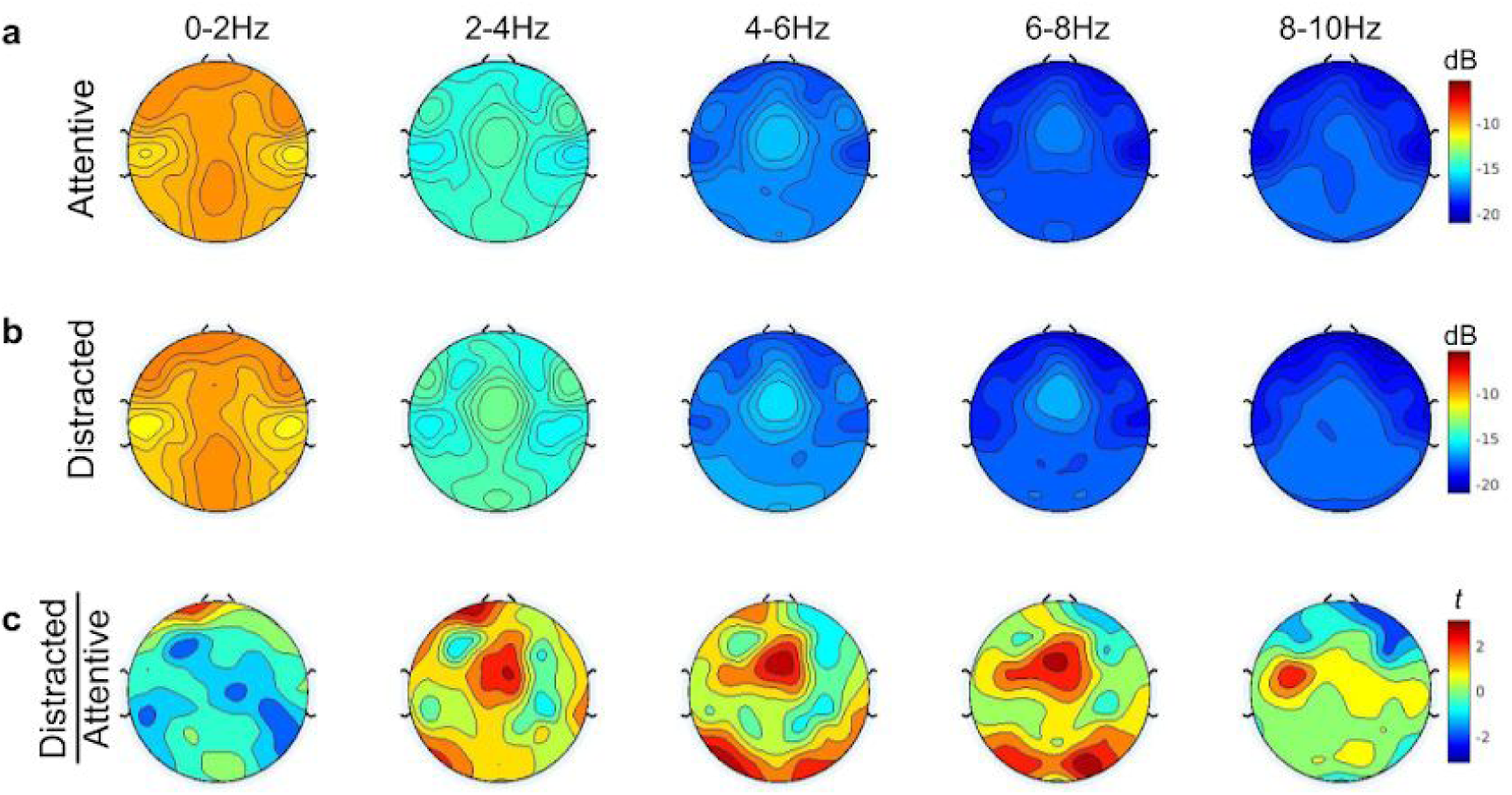
Working Memory Paradigm Validation. The figure depicts topological normalized average power of 0-2Hz, 2-4Hz, 4-6Hz, 6-8Hz, and 8-10Hz bands. during the (a) low-load attentive condition, and, (b) high-load distracted condition (b), with, (c) a channel pairwise t-test t’s representing the distracted over the attentive condition.

**Figure S3.**
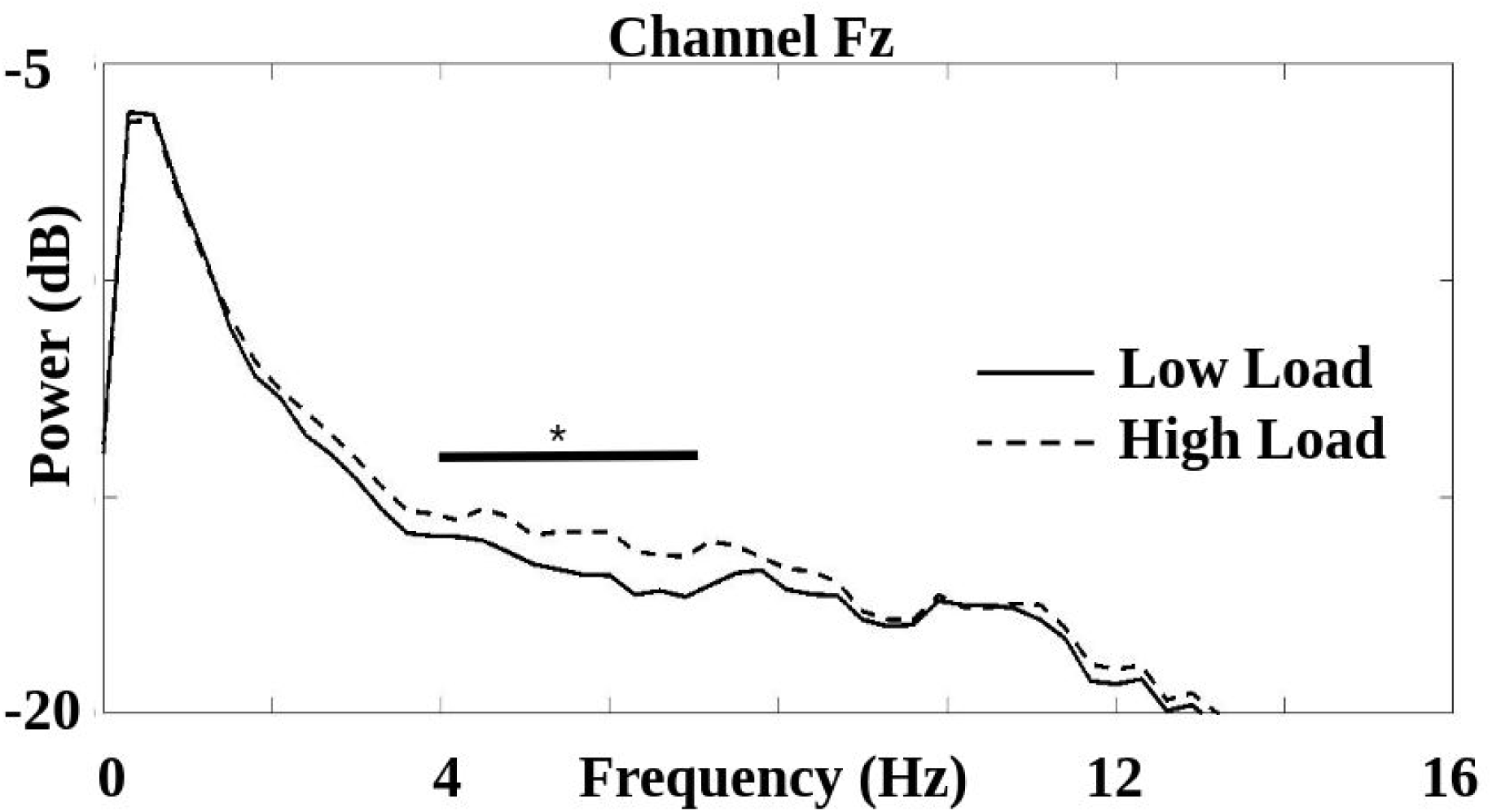
Power (dB) over frequency (Hz) for channel Fz during high and low cognitive load. *; *p*<0.05

